# Distinct mesoderm migration phenotypes in extra-embryonic and embryonic regions of the early mouse embryo

**DOI:** 10.1101/438721

**Authors:** Bechara Saykali, Navrita Mathiah, Wallis Nahaboo, Marie-Lucie Racu, Matthieu Defrance, Isabelle Migeotte

## Abstract

In the gastrulating mouse embryo, epiblast cells delaminate at the primitive streak to form mesoderm and definitive endoderm, through an epithelial-mesenchymal transition.

Mosaic expression of a membrane reporter in nascent mesoderm enabled recording cell shape and trajectory through live imaging. Upon leaving the streak, cells changed shape and extended protrusions of distinct size and abundance depending on the neighboring germ layer, as well as the region of the embryo. Embryonic trajectories were meandrous but directional, while extra-embryonic mesoderm cells showed little net displacement.

Embryonic and extra-embryonic mesoderm transcriptomes highlighted distinct guidance, cytoskeleton, adhesion, and extracellular matrix signatures. Specifically, intermediate filaments were highly expressed in extra-embryonic mesoderm, while live imaging for F-actin showed abundance of actin filaments in embryonic mesoderm only. Accordingly, *RhoA* or *Rac1* conditional deletion in mesoderm inhibited embryonic, but not extra-embryonic mesoderm migration.

Overall, this indicates separate cytoskeleton regulation coordinating the morphology and migration of mesoderm subpopulations.

## INTRODUCTION

In mice, a first separation of embryonic and extra-embryonic lineages begins in the blastocyst at embryonic day (E) 3.5 when the trophectoderm is set aside from the inner cell mass. A second step is the segregation of the inner cell mass into the epiblast, the precursor of most fetal cell lineages, and the extra-embryonic primitive endoderm (Chazaud & Yamanaka, 2016). At E6, the embryo is cup-shaped and its anterior-posterior axis is defined. It comprises three cell types, arranged in two layers: the inner layer is formed by epiblast, distally, and extra-embryonic ectoderm, proximally; the outer layer, visceral endoderm, covers the entire embryo surface. The primitive streak, site of gastrulation, is formed at E6.25 in the posterior epiblast, at the junction between embryonic and extra-embryonic regions, and subsequently elongates to the distal tip of the embryo. The primitive streak is the region of the embryo where epiblast cells delaminate through epithelial-mesenchymal transition to generate a new population of mesenchymal cells that form the mesoderm and definitive endoderm layers.

All mesoderm, including the extra-embryonic mesoderm, is of embryonic epiblast origin. At the onset of gastrulation, emerging mesoderm migrates either anteriorly as so-called embryonic mesodermal wings, or proximally as extra-embryonic mesoderm (Arnold & Robertson, 2009; Sutherland, 2015). Cell lineages studies showed that there is little correlation between the position of mesoderm progenitors in the epiblast and the final localization of mesoderm descendants (Lawson, Meneses, & Pedersen, 1991). Rather, the distribution of mesoderm subpopulations depends on the temporal order and anterior-posterior location of cell recruitment through the primitive streak (Kinder et al., 1999). Posterior primitive streak cells are the major source of extra-embryonic mesoderm, while cells from middle and anterior primitive streak are mostly destined to the embryo proper. However, there is overlap of fates between cells delaminating at different sites and timings (Kinder et al., 1999, 2001). Extra-embryonic mesoderm contributes to the amnion, allantois, chorion, and visceral yolk sac. It has important functions in maternal-fetal protection and communication, as well as in primitive erythropoiesis (Watson, 2005). Embryonic mesoderm separates into lateral plate, intermediate, paraxial and axial mesoderm, and ultimately gives rise to cranial and cardiac mesenchyme, blood vessels and hematopoietic progenitors, urogenital system, muscles and bones, among others. Endoderm precursors co-migrate with mesoderm progenitors in the wings and undergo a mesenchymal-epithelial transition to intercalate into the visceral endoderm (Viotti, Foley, & Hadjantonakis, 2014).

Mesoderm migration mechanisms have mostly been studied in fly, fish, frog and chicken embryos. Interestingly, during fly gastrulation, mesodermal cells migrate as a collective (Bae, Trisnadi, Kadam, & Stathopoulos, 2012). In the fish prechordal plate, all cells have similar migration properties but they require contact between each other for directional migration (Dumortier, Martin, Meyer, Rosa, & David, 2012). Live imaging experiments in chick suggest a collective mode of migration, even though cells frequently make and break contacts with their neighbors (Chuai, Hughes, & J. Weijer, 2012).

Relatively little is known about mesoderm migration in mice because most mutant phenotypes with mesodermal defects result from anomalies in primitive streak formation, mesoderm specification, or epithelial-mesenchymal transition (Arnold & Robertson, 2009), precluding further insight into cell migration mechanisms. We previously identified a role for the Rho GTPase Rac1, a mediator of cytoskeletal reorganization, in mesoderm migration and adhesion (Migeotte, Grego-Bessa, & Anderson, 2011).

Recent advances in mouse embryo culture and live imaging have overcome the challenge of maintaining adequate embryo growth and morphology while performing high-resolution imaging. It facilitated the uncovering of the precise spatial and temporal regulation of cellular processes and disclosed that inaccurate conclusions had sometimes been drawn from static analyses (Viotti et al., 2014). Live imaging of mouse embryos bearing a reporter for nuclei has pointed towards individual rather than collective migration in the mesodermal wings (Ichikawa et al., 2013). Very recently, a spectacular adaptive light sheet imaging approach allowed reconstructing fate maps at the single cell level from gastrulation to early organogenesis (McDole et al., 2018). However, little is known about how mesoderm populations regulate their shape and migration mechanisms as they travel across distinct embryo regions to fulfill their respective fates.

Here, high-resolution live imaging of nascent mesoderm expressing membrane-bound GFP was used to define the dynamics of mesoderm cell morphology and its trajectories. Mesoderm cells exhibited a variety of cell shape changes determined by their spatial localization in the embryo, and the germ layer they were in contact with. The embryonic mesoderm migration path was meandrous but directional, and depended on the Rho GTPases RhoA and Rac1. Extra-embryonic mesoderm movement was, strikingly, GTPases independent. Transcriptomes of different mesoderm populations uncovered specific sets of guidance, adhesion, cytoskeleton and matrix components, which may underlie the remarkable differences in cell behavior between mesoderm subtypes.

## RESULTS

### Mesoderm migration mode and cell shape differ in embryonic versus extra-embryonic regions

The T box transcription factor *Brachyury* is expressed in posterior epiblast cells that form the primitive streak, maintained in cells that delaminate through the streak, then down-regulated once cells progress anteriorly in the mesodermal wings (Wilkinson, Bhatt, & Herrmann, 1990). In order to visualize nascent mesoderm, *T*(s)::Cre (hereafter referred to as *T*-Cre) transgenic animals, in which Cre expression is driven by the regulatory elements of the *Brachyury* gene (Feller, Schneider, Schuster-Gossler, & Gossler, 2008; Stott, Kispert, & Herrmann, 1993), were crossed to a membrane reporter line: Rosa26::membrane dtTomato/membrane GFP (Muzumdar, Tasic, Miyamichi, Li, & Luo, 2007) (referred to as mTmG) (Fig. 1). In*T*-Cre; mTmG embryos, primitive streak and mesoderm derived cells have green membranes (mG), whereas all other cells have red membranes (mG). Embryos dissected at E6.75 or E7.25 were staged according to (K M Downs & Davies, 1993) (Sup. Fig. 1a) and examined in different orientations by confocal or two-photon excitation live imaging for 8 to 12 hours (Fig. 1, Sup. Fig. 1b and c, Videos 1 and 2). Conversion of mT to mG was first observed at Early/Mid Streak (E/MS) stage, and was initially mosaic, which facilitated the tracking of individual migrating cells with high cell shape resolution. From Mid/Late Streak (M/LS) onwards, most primitive streak cells underwent red to green conversion (Sup. Fig. 1d, e).

**Figure 1:**
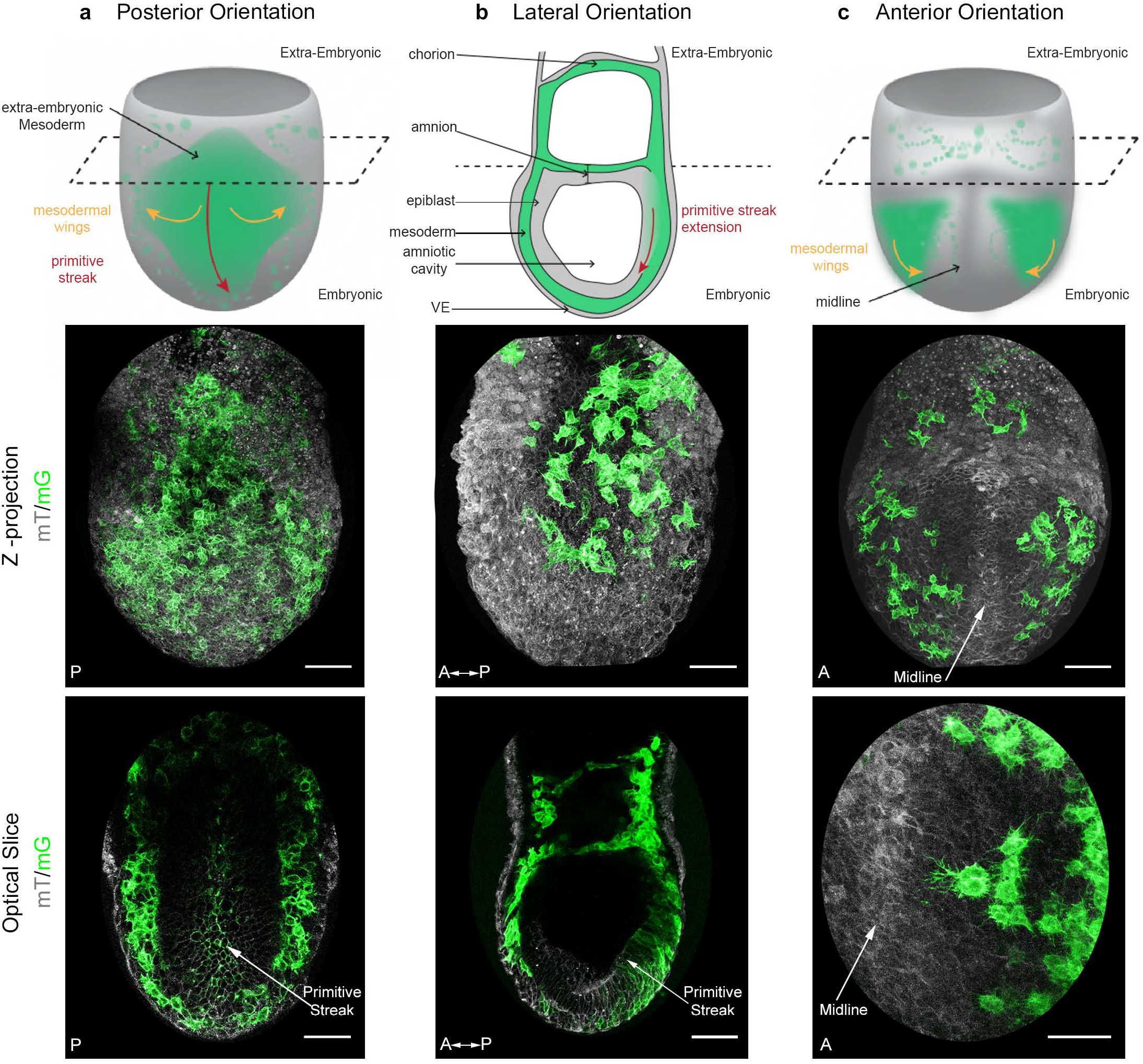
Mosaic membrane GFP labeling of nascent mesoderm allows following individual cell migration through embryo live imaging. (a) Posterior view. Top: 3D scheme with mesoderm layer in green and the rest of the embryo in grey. The dashed line separates embryonic and extra-embryonic regions. Middle: Z-projection of two-photon stack. Bottom: optical slice highlighting the primitive streak. (b) Lateral view, anterior to the left. Top: 2D scheme. Middle: Z-projection of two-photon stack showing cells progression from posterior to anterior. Bottom: sagittal optical slice. (c) Anterior view. Top: 3D scheme. Middle: Z-projection of two-photon stack with most anterior cells reaching the midline. Bottom: optical slice zoomed on filopodia extending towards the midline. All embryos were dissected at E7.25 and are at Late Streak/Zero Bud stage.VE: Visceral Endoderm; mG: membrane GFP, in green; mT: membrane dtTomato, in grey. (Scale bars: 50 μm).

The shape of mesoderm cells and their tracks were recorded through imaging of embryos from different perspectives between ES and Early Bud (EB) stages of development, in order to obtain images of optimal quality for each embryo region (Fig. 1 and Videos 1 and 2). Posterior views (Fig. 1a) showed proximal to distal primitive streak extension and rounding of cells exiting the streak. Lateral views (Fig. 1b allowed comparing cells as they migrated laterally in mesodermal wings, or proximally in extra-embryonic region. Anterior views (Fig. 1c) showed cell movement towards the midline. The imaging time frame did not allow following individual cells from their exit at the primitive streak to their final destination. However, the trajectories we acquired fitted with the fate maps built using cellular labeling or transplantation (Kinder et al., 1999, 2001), or adaptive light sheet microscopy (McDole et al., 2018) (Videos 3 and 4). The first converted (GFP positive) cells in ES embryos dissected around E6.75 usually left the posterior site of the primitive streak to migrate towards the extra-embryonic compartment. Embryonic migration started almost simultaneously, and migration towards both regions proceeded continuously.

Strikingly, migration behavior (Fig. 2a, Videos 3 and 4) and cell shape (Fig. 2b) were different depending on the region cells migrated into. In the embryonic region, mesoderm cells had a global posterior to anterior path, even though they zigzagged in all directions (proximal-distal, left-right, and even anterior-posterior). Cells did not migrate continuously, but showed alternations of tumbling behavior with straighter displacement. Embryonic mesoderm cells from ES/MS embryos tracked for 2.5 h moved at a mean speed of 0.65 μm/min to cover approximately 90 μm and travel a net distance of 40 μm (Fig. 2a’, Table 1). Straightness (the ratio of net over travel displacement, so that a value of 1 represents a linear path) was approximately 0.5 (Fig. 2a’, Table 1). Cells in the extra-embryonic region moved slightly slower (0.45 μm/min) to do approximately 70 μm, but their net displacement (20 μm) and straightness (0.3) were significantly smaller, reflecting trajectories with no obvious directionality (Fig. 2a’, Table 1, and Video 5).

**Figure 2:**
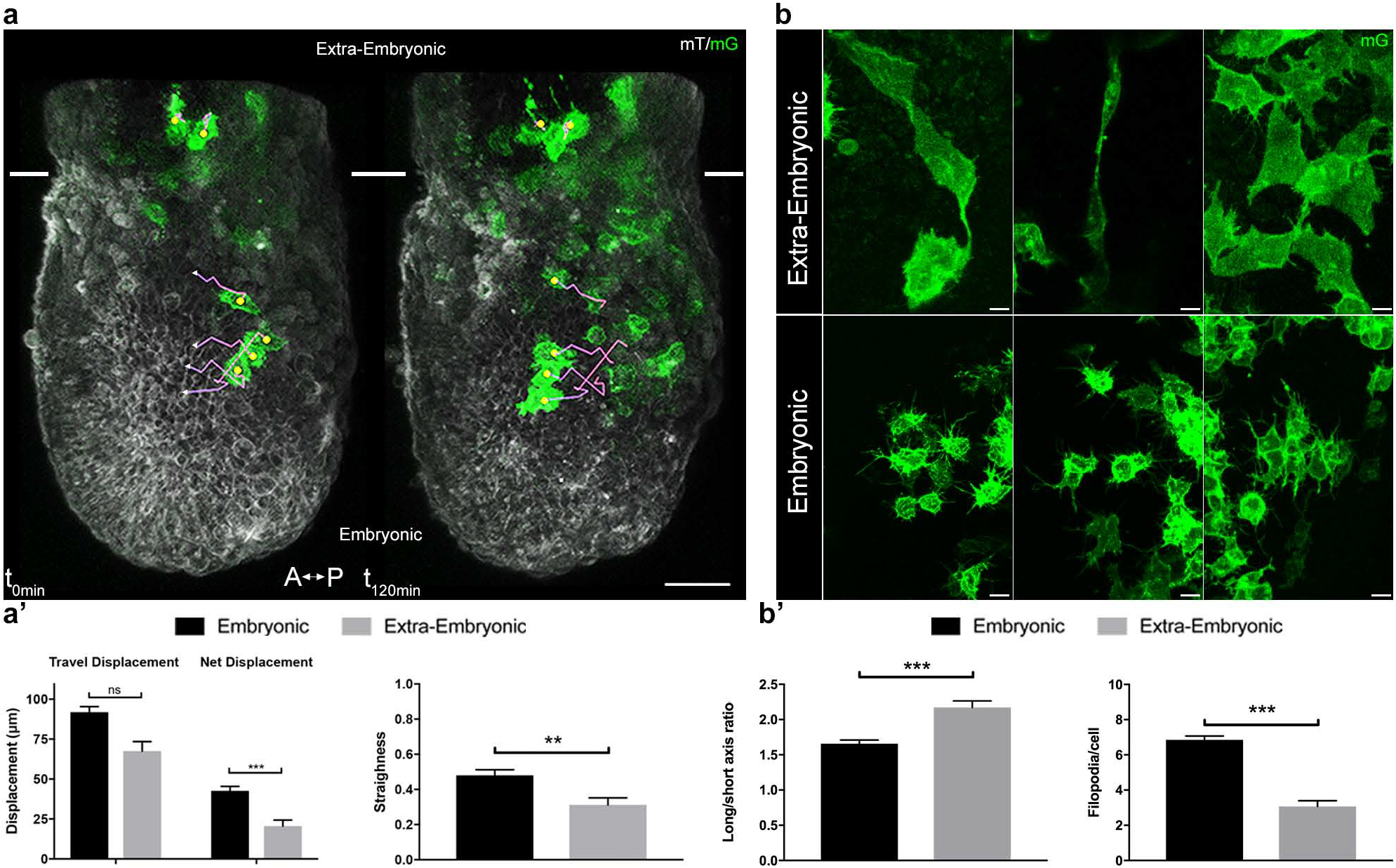
Embryonic and extra-embryonic mesoderm populations have different morphology and migration pattern. (a) Z-projections of confocal stacks from a *T*-cre; mTmG embryo dissected at E6.75 (Early Streak), with cell migration tracking for 120 min. Anterior to the left. White lines mark the embryonic/extra-embryonic boundary. (Scale bar: 50 μm). (a’) Quantification (mean ± SEM) of travel and net displacement (Left) and path straightness (Right, on a scale of Oto 1) of embryonic (black, n=34 from 4 Early/Mid Streak embryos) and extra-embryonic (grey, n= 17 cells) mesoderm cells. Data can be found in Table 1 and Source Data 1. (b) Embryonic and extra-embryonic mesoderm cell shapes (images extracted from 4 Late Streak embryos) (Z-projections of two-photon stacks, scale bar: 10 μm). (b’) Left: Long/short axis ratio of 2D inner ellipse as quantification of cell stretch (mean± SEM, n= 85 embryonic cells in black, n= 83 extra-embryonic cells in grey, out of 8 Mid Streak to Zero Bud stages embryos). Right: Quantification (mean ± SEM) of number of filopodia per cell in embryonic (black, n= 167 cells out of 5 Mid Streak to Early Bud stages embryos) and extra-embryonic (grey, n=28 cells) mesoderm cells. Data can be found in Table 2, Source Data 2 and 3. *P* values were calculated using the Mann-Whitney-Wilcoxon. mG: membrane GFP, in green; mT: membrane dtTomato, in grey.

**Table 1:** Tracking details for embryonic and extra-embryonic mesoderm. Cells were tracked for approximately 150 min. *P* values were calculated using the Mann– Whitney– Wilcoxon. Data can be found in Source Data 1.

|  | Net displacement (μm) |  | Travel displacement (μm) |  | Straightness |  | Mean speed (μm/min) |  |  |
| --- | --- | --- | --- | --- | --- | --- | --- | --- | --- |
|  | Mean | SEM | Mean | SEM | Mean | SEM | Mean | SEM | N |
| Extra-embryonic | 20,58 | 3,74 | 67,50 | 6,01 | 0,31 | 0,04 | 0,44 | 0,03 | 17 |
| Embryonic | 42,64 | 2,79 | 91,86 | 3,51 | 0,48 | 0,03 | 0,67 | 0,03 | 34 |
| P-value | 7,93E-05 |  | 5,29E-01 |  | 2,48E-03 |  | 2,00E-06 |  |  |

Extra-embryonic cells were stretched, sometimes spanning the entire width of the embryo, and twice larger (Fig. 2b, b’, Table 2). They had few large protrusions, and filopodia were scarce and short (Fig. 2b, b’, Table 2).

**Table 2:**
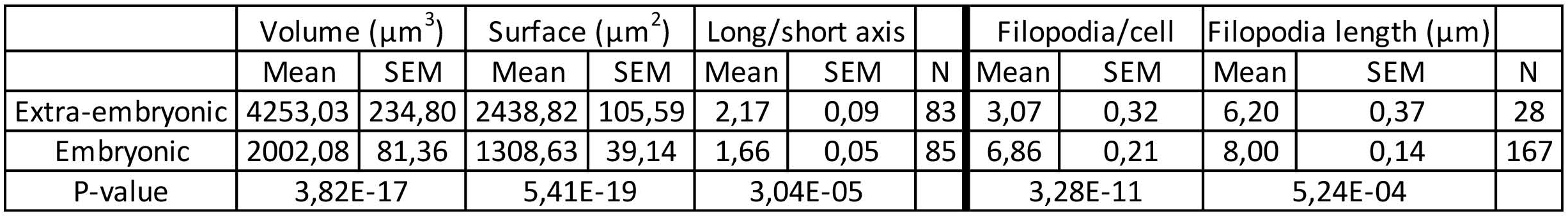
Cell shape, size, and filopodia comparison between embryonic and extra-embryonic mesoderm. *P* values were calculated using the Mann–Whitney–Wilcoxon for surface, long/short axis and filopodia/cell, and the *t* test for volume, and filopodia length. Data can be found in Source Data 2 and 3.

### Mesoderm cells have distinct morphology depending on their interaction with different germ layers

Cells passing through the primitive streak were, as reported (Ramkumar et al., 2016; M. Williams, Burdsal, Periasamy, Lewandoski, & Sutherland, 2012), bottle shaped with a basal round cell body and an apical thin projection (Fig. 3a). Mesoderm cells in contact with the epiblast and visceral endoderm sent thin protrusions towards their respective basal membranes (Fig. 3b, b’, and Video 6). Interestingly, the density of thin protrusions was much higher in cells in contact with the visceral endoderm; this phenotype was observed as early as ES, while intercalation of prospective definitive endoderm cells in the visceral endoderm layer occurred only from LS stage onwards (Viotti et al., 2014). Cells in tight clusters surrounded by other mesoderm cells in the wings had a smoother contour. A caveat was that protrusions couldn’t be visualized between two cells of similar membrane color. Reconstruction of cells in the anterior part of the wings, where recombination was incomplete, showed thin protrusions between mesoderm cells. Cells also extended long broad projections, which spanned several cell diameters and were sent in multiple directions before translation of the cell body, in what seemed a trial and error process (Fig. 3c and Videos 1, 2 and 7). The presence of potential leader cells could not be assessed, as the first cells converted to green are not the most anterior ones (Sup. Fig. 1d). Nonetheless, cells with scanning behavior were observed at all times, which suggests that all cells are capable to explore their surroundings.

**Figure 3:**
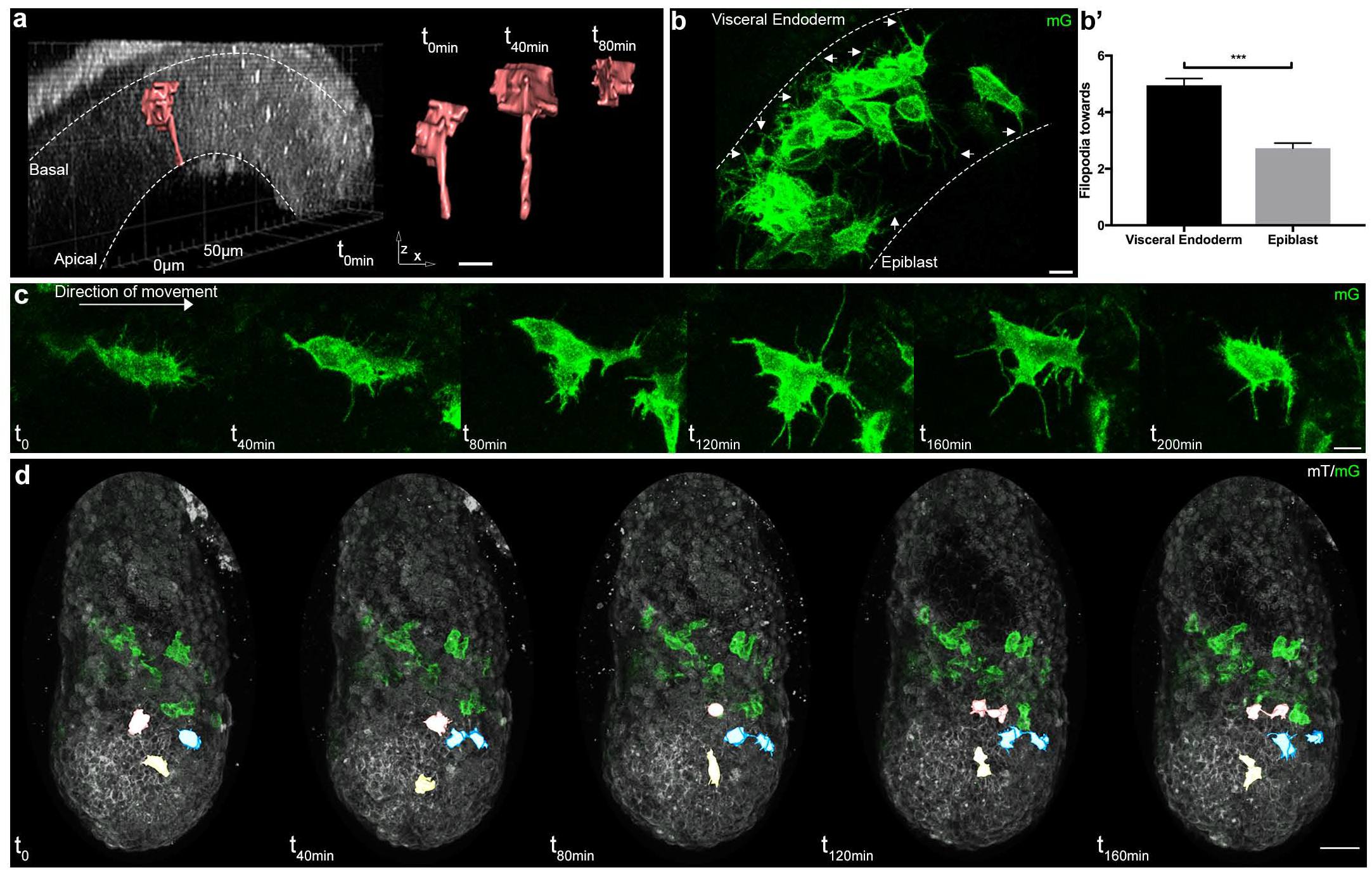
Cell shape changes of migrating mesoderm. (a) Cell shape progression of nascent mesoderm delaminating at the primitive streak of a Mid Streak embryo (Z-projection of two-photon stack, scale bar: 10 μm). (b) Mesoderm cells extend filopodia (arrows) towards epiblast and visceral endoderm. Embryo is at Late Streak stage. (Z-projection of two-photon stack, scale bar: 10 μm). (b’) Quantification of filopodia (filopodia per cell, mean± SEM, n=40 cells out of 4 Late Streak embryos for each, p<0.0001). *P* value was calculated using the *t* test. Data can be found in Source Data 4. (c) Montage of a mesoderm cell (from a Mid Streak stage embryo) displaying seeking behavior (Z-projection of two-photon stack, scale bar: 10 μm). (d) Mesoderm cells are highlighted, through manual segmentation, in red, blue and yellow to track cell behavior after division (Z-projection of confocal stacks from a Mid Streak stage embryo, anterior view, scale bar: 50 μm). mG: membrane GFP, in green; mT: membrane dtTomato, in grey.

### Cell-cell communication within the mesoderm layer

As expected from fate mapping experiments, daughter cells resulting from mitosis within the mesoderm layer followed close and parallel trajectories (Fig. 3d and Source Data 5). They travelled a similar net distance over 204 min (net displacement ratio: 0.91 ± 0.01, n=12 pairs from four embryos at E/MS stage), in the same direction (angle: 7 ± 1.13°), with one daughter cell displaying a higher straightness (travel displacement ratio: 0.61 ± 0.07). They remained close to one another (mean distance between daughter cells: 15.6 ± 2.35 μm), but not directly apposed. Interestingly, they stayed linked by thin projections for hours, even when separated by other cells (Fig. 3d).

Mesoderm cell migration is often compared to neural crest migration, as both cell types arise through epithelial-mesenchymal transition (Roycroft & Mayor, 2016). An important feature of neural crest migration is contact inhibition of locomotion, where cells that collide tend to move in opposite directions. In contrast, most mesoderm cells stayed in contact upon collision (Source Data 6): 16 out of 24 cell pairs from 5 ES to Zero Bud (0B) embryos remained attached for 2.5 hour (one briefly lost contact before re-joining), 3/24 stayed joined for around 1 hour, and 5/24 pairs separated instantly. We segmented 8 pairs for 166 min, and observed a mean distance at the end of tracking of 52.5 ± 32.5 μm; they followed parallel trajectories (angle: 8.25 ± 1.7°, n=8 pairs), for a similar net distance (net displacement ratio: 0.85 ± 0.07; travel displacement ratio: 0.64 ± 0.11). Thin projections could occasionally be observed between them after contact.

### Embryonic and extra-embryonic mesoderm molecular signatures

Embryonic and extra-embryonic mesoderm cells were isolated through fluorescence (GFP)-assisted cell sorting from E7.5 MS and LS *T*-Cre; mT/mG embryos in order to generate transcriptomes. Biological replicates (4 MS, 2 LS) from both stages resulted in grouping of samples according to the embryonic region (Fig. 4a, b). Non-supervised clustering based on embryonic and extra-embryonic gene signatures identified by single cell sequencing in (Scialdone et al., 2016) showed that samples segregated as expected (not shown). We performed pairwise comparison of embryonic and extra-embryonic samples obtained at both stages and selected the genes that were consistently differentially expressed with a fold change > 2. Gene ontology analysis showed significant enrichment in expected cellular (transcription and differentiation), developmental (angiogenesis, heart development, somitogenesis, and hematopoiesis), and signaling (Wnt, BMP and Notch) biological processes. Interestingly, differences were also seen in gene clusters involved in migration, adhesion, matrix organization, and small GTPases mediated signal transduction (Sup. Fig. 2a).

**Figure 4:**
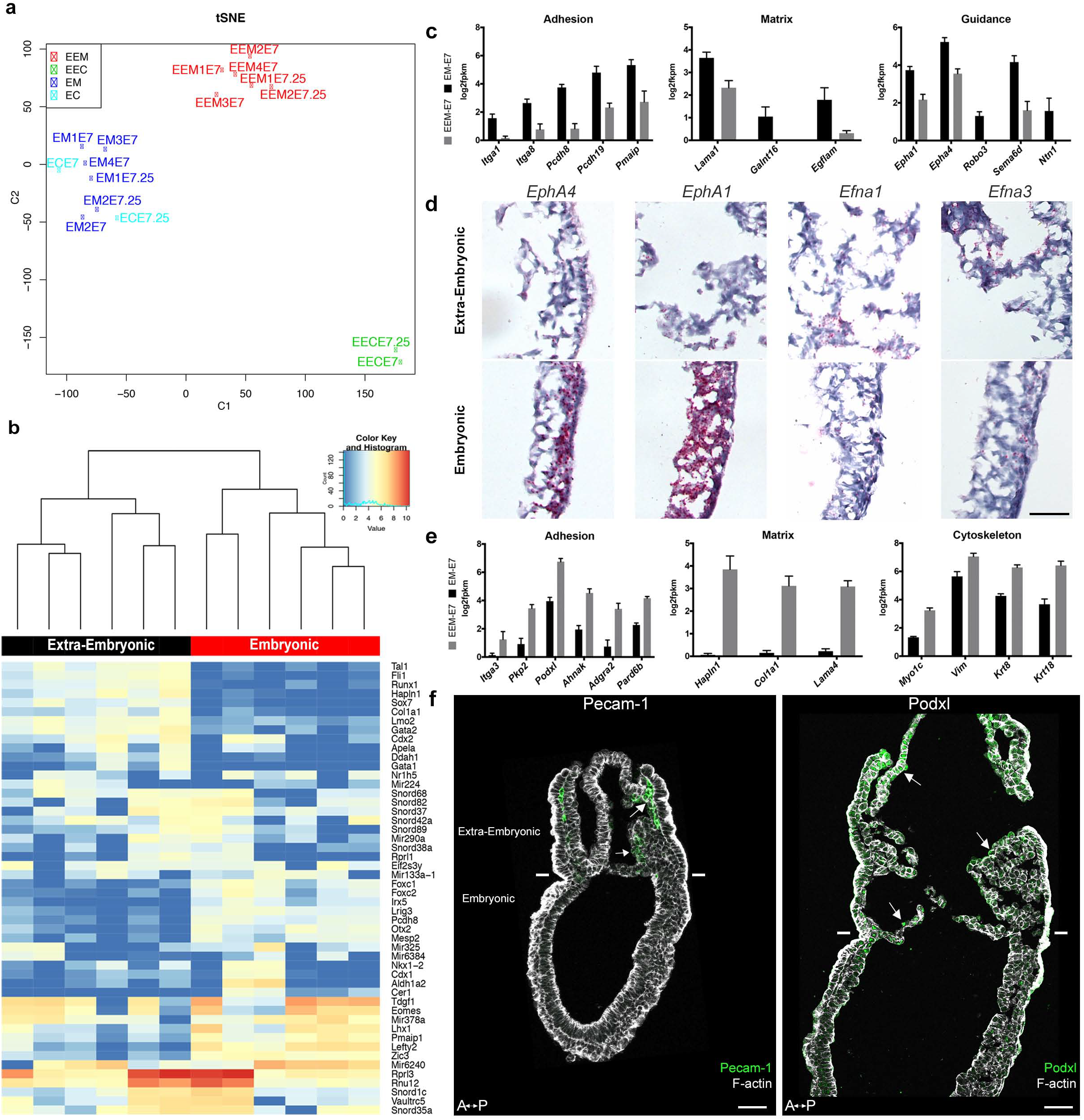
Transcriptomes of mesoderm populations identify differences between embryonic and extra-embryonic mesoderm. (a) t-Distributed Stochastic Neighbor Embedding (t-SNE) confirms grouping of similar biological samples at E? (Mid Streak stage) and E?.25 (Late Streak stage). EM: Embryonic mesoderm; EEM: Extra-Embryonic mesoderm; EC: Embryonic control; EEC: Extra-Embryonic control. More sample information can be found in Source Data 7. (b) Heat map showing differentially expressed genes between embryonic and extra-embryonic mesoderm with the highest statistical significance. (c, e) Selection of genes with higher expression in embryonic (c) or extra-embryonic (e) mesoderm, represented as mean ± SEM of log2 fpkm at E? (n=4 biological replicates, p<0.01), with embryonic in black and extra-embryonic in grey. All represented genes are also significantly differentially regulated at E?.25. Data can be found in Source Data 8 and 9. (d) *In situ* hybridization of sagittal sections (anterior to the left) from Zero to Early Bud stages embryos highlighting transcripts for *EphA4, EphA1, Efna1* and *Efna3,* represented by red dots, in the posterior region. Entire embryo sections are shown in Sup. Fig. 2. (Scale bar: 100 μm) (f) Sagittal sections (anterior to the left) from Early Bud stage embryos stained for Platelet endothelial cell adhesion molecule 1 (left panel, Pecam-1 in green, Z-projection of confocal stack), Podocalyxin (right panel, Podxl in green, optical slice), and F-actin (Phalloidin, grey). See same section stained for mGFP in Sup. Fig. 2. White lines mark the embryonic/extra-embryonic boundary. (Scale bars: 50 μm).

Genes with known expression pattern in gastrulating embryos found enriched in embryonic mesoderm included well-described transcription factors, as well as FGF, Wnt, Notch, TGFβ and Retinoic Acid effectors (Sup. Fig. 2b). Genes expected to be more expressed in extra-embryonic mesoderm included the transcription factors *Ets1* and *Tbx20*, and several members of the TGFβ pathway (Inman & Downs, 2007; Pereira et al., 2011) (Sup. Fig. 2c). Primitive hematopoiesis, the initial wave of blood cell production which gives rise to primitive erythrocytes, macrophages, and megakaryocytes, takes place around E7.25 in hemogenic angioblasts of the blood islands (Lacaud & Kouskoff, 2017). Expression of genes involved in hemangioblast development, endothelium differentiation, and hematopoiesis increased from MS to LS (Fig. 4f and Sup. Fig. 2c). In addition, we confirmed two extra-embryonic genes identified through subtractive hybridization at E7.5 (Kingsley et al., 2001): *Ahnak* (Sup. Fig. 2c, see also (Karen M. Downs, McHugh, Copp, & Shtivelman, 2002)) and the imprinted gene *H19* (Sup. Fig. 2c).

Of particular interest among the genes with higher expression in embryonic mesoderm for which no expression data was available at the stage of development were genes related to matrix (*Lama1, Galnt16, Egflam*), adhesion (*Itga1, Itga8, Pcdh8, Pcdh19, Pmaip1*), and guidance (*Epha1* and *4*, *Robo3*, *Sema6d, Ntn1*) (Fig. 4c). *EphA4* expression in the mouse embryo has been described in the trunk mesoderm and developing hindbrain at Neural Plate (NP) stage (M A Nieto, Gilardi-Hebenstreit, Charnay, & Wilkinson, 1992). In LS embryos, *Epha4* expression was higher in the primitive streak and embryonic mesoderm (Fig. 4d and Sup. Fig. 2e). Dynamic *Epha1, Efna1 and Efna3* expression patterns have been shown during gastrulation (Duffy, Steiner, Tam, & Boyd, 2006). In LS/0B embryos, *Epha1* mRNA was present in the primitive streak, mostly in its distal part. Its ligand *Efna1* was in the primitive streak with an inverse gradient, and was mainly expressed in the extra-embryonic region, notably in amnion and in chorion. *Efna3* was very abundant in the chorion (Fig. 4d and Sup. Fig. 2e).

In parallel, in extra-embryonic mesoderm, we found higher expression of distinct sets of genes with putative roles in guidance (*Unc5c, Dlk1, Scube2, Fzd4*), matrix composition (*Hapln1, Col1a1, Lama4*), adhesion (*Itga3, Pkp2, Podxl, Ahnak, Adgra2, Pard6b*), Rho GTPase regulation (*Rasip1, Stard8, RhoJ*), and cytoskeleton (*Myo1c, Vim*, *Krt8* and *Krt18)* (Fig. 4e and not shown). Interestingly, Podocalyxin (*Podxl)* was abundant in extra-embryonic mesoderm (Fig. 4f and Sup. Fig. 2d), which fits with data from embryo and embryoid body single cell sequencing showing that *Podxl* is a marker for early extra-embryonic mesoderm and primitive erythroid progenitors of the yolk sac (Zhang et al., 2014).

### Embryonic and extra-embryonic mesoderm cells have distinct cytoskeletal composition

In view of the differences in cell shape and migration, we focused on the cytoskeleton, in particular actin, and intermediate filaments proteins (Vimentin and Keratins). Vimentin was found in all mesoderm cells as expected, but more abundant in extra-embryonic mesoderm (Fig. 5a). Remarkably, within the mesoderm layer, Keratin 8 was selectively expressed in extra-embryonic mesoderm cells (amniochorionic fold, amnion, chorion, and developing allantois) (Fig. 5b-d). In contrast, the filamentous actin (F-actin) network, visualized by Phalloidin staining, seemed denser in embryonic mesoderm (Fig. 5a). To visualize F-actin only in mesoderm, we took advantage of a conditionally inducible mouse model expressing Lifeact-GFP, a peptide that binds specifically to F-actin with low affinity, and thus reports actin dynamics without disrupting them (Schachtner et al., 2012). Live imaging of *T*-Cre; LifeAct-GFP embryos at MS and LS stage confirmed that while LifeAct-GFP positive filaments could be visualized clearly in embryonic mesoderm cells, GFP was weaker and diffuse in extra-embryonic mesoderm (Fig. 5e and Video 8).

**Figure 5:**
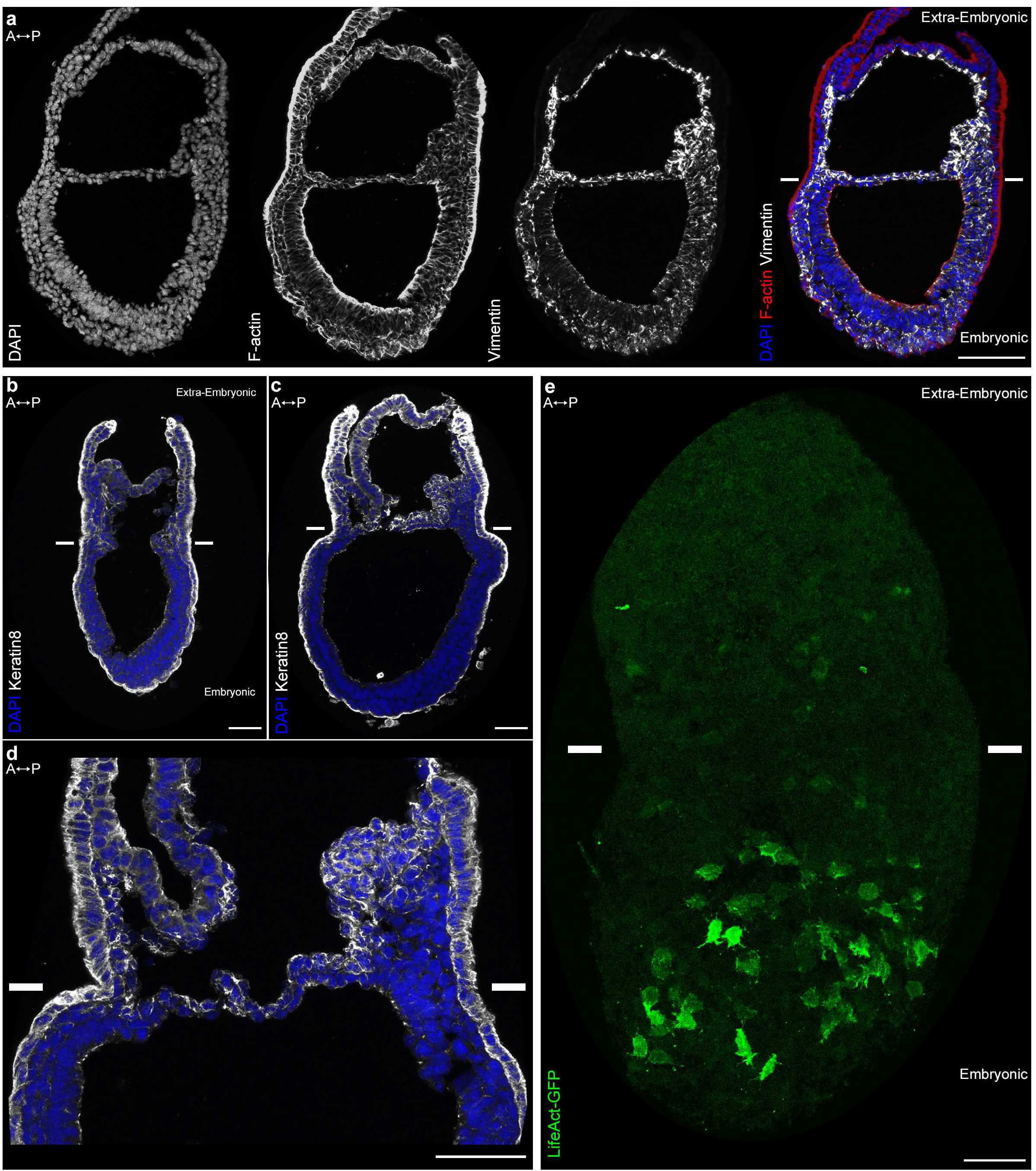
Embryonic and extra-embryonic mesoderm cells have distinct cytoskeleton composition. (a) Z-projections of confocal stack of a sagittal section from an Early Bud stage embryo stained for Vimentin, F-Actin (Phalloidin), and nuclei (DAPI). (b, d) Z-projections of confocal stacks of sagittal sections from Late Streak (b) and Early Bud (c: 20x, d: 40x) stages embryos stained for Keratin 8 and nuclei (DAPI). (e) Z-projection of two-photon stack of a whole-mount *T-Cre;* LifeAct-GFP Late Streak embryo. Anterior is to the left. (Scale bars: 50 μm)

### Extra-embryonic mesoderm migration is Rho GTPases independent

Rho GTPases are molecular switches that relay signals from cell surface receptors to intracellular effectors, leading to a change in cell behavior (Hodge & Ridley, 2016). They are major regulators of cytoskeletal rearrangements (Hall, 1998), and the spatiotemporal fine regulation of Rho GTPases activities determines cytoskeletal dynamics at the subcellular level (Spiering & Hodgson, 2011). Therefore, inactivation of a given Rho GTPase may result in variable consequences depending on cell type and context. We previously established that *Sox2*-Cre mediated deletion of *Rac1* in the epiblast before onset of gastrulation causes impaired migration of embryonic mesoderm while extra-embryonic mesoderm migration is less severely affected (Migeotte et al., 2011). We thus hypothesized that Rho GTPases might be differentially regulated in cells invading both regions, resulting in some of the observed distinctions in cytoskeletal dynamics, cell shape and displacement mode.

Mutations were induced in cells transiting the primitive streak by crossing heterozygous wild-type/null *RhoA* (Jackson et al., 2011) or *Rac1* (Walmsley, 2003) animals bearing the *T*-Cre transgene with animals homozygous for their respective conditional alleles bearing the mTmG reporter (mutant embryos are referred to as *RhoAΔmesoderm and Rac1Δmesoderm)*. The phenotypes of *RhoAΔmesoderm and Rac1Δmesoderm embryos* were less severe than that of *RhoA^Δepi^ and Rac1^Δepi^* embryos (our unpublished data and (Migeotte et al., 2011)) (Sup. Fig. 3). Mutants were morphologically indistinguishable at E7.5. At E8.5, *RhoAΔmesoderm* embryos were identified, though with incomplete penetrance, as being slightly smaller than their wild-type littermates (11/12 mutants had a subtle phenotype, including 5 with reduced numbers of somites, and 6 with abnormal heart morphology) (Sup. Fig. 3a). By E9.5, all *RhoAΔmesoderm* embryos had an obvious phenotype (12/12 mutants had a small heart, 9/12 had a reduced number of somites, 2/12 had an open neural tube, 2/12 had a non-fused allantois) (Sup. Fig. 3b). *Rac1Δmesoderm* embryos also had subtle phenotypes at E8.5 (15/16 embryos were slightly smaller than wild-type littermates, 4/16 had a small heart) (Sup. Fig. 3d). *In situ* hybridization for *Brachyury* showed weaker staining in the tail region in 5/10 mutant embryos, indicative of reduced presomitic mesoderm (Sup. Fig. 3c). By E9.5, all mutants had abnormal heart morphology and reduced body length, and 3 embryos out of 9 were severely delayed (Sup. Fig. 3e). At E10.5, penetrance was complete; 7/7 embryos had a short dysmorphic body and pericardial edema (not shown). The phenotypic variability at early time points likely reflects mosaicism of *T*-Cre mediated recombination.

Embryonic mesoderm explants from E7.5 MS/LS mTmG; *Rac1Δmesoderm* or *RhoAΔmesoderm* embryos were plated on fibronectin. In wild-type explants, cells showed a radial outgrowth of the explants, displaying large lamellipodia in the direction of migration (Video 9). After cell-cell contact, they remained connected through long thin filaments. *RhoA* deficient explants showed no outgrowth of individual cells (Sup. Fig. 4a). *RhoA* mutant cells appeared more cohesive and displayed fewer protrusions than wild-type cells. Remarkably, live imaging of *RhoA* deleted explants showed a phase of compaction preceding cell migration (Video 10; 2/4 *RhoAΔmesoderm* mutant explants displayed compaction). In *Rac1* explants (Sup. Fig. 4b), GFP-expressing cells remained within the domain of the dissected explant and displayed pycnotic nuclei, while wild-type non-GFP cells could migrate. Live imaging could not be performed as 4 out of 5 mutant explants detached from the plate. This is similar to explants from *Rac1* epiblast-specific mutants (Migeotte et al., 2011), and is attributed to lack of adhesion-dependent survival signals.

Live imaging of mTmG; *RhoAΔmesoderm or Rac1Δmesoderm* embryos dissected at E6.75 or 7.25 (Fig. 6) showed that the majority of *RhoA* and *Rac1* mesoderm-specific mutants (4/8 for *RhoA*, 6/9 for *Rac1*) displayed an accumulation of cells at the primitive streak, which formed a clump on the posterior side between epiblast and visceral endoderm (Fig. 6a, b and Video 11), indicating a mesoderm migration defect. Interestingly, although embryonic mesoderm migration was impaired, with only a handful of cells visible on the anterior side by E7.5, extra-embryonic mesoderm migration was maintained (Fig. 6a’, b’ and Videos 12, 13). Accordingly, staining for mesoderm-derived vascular structures (Pecam-1) in the yolk sac at E8.5 showed no difference between mutants and wild-type embryos (Sup. Fig. 4c, d). Those findings suggest that extra-embryonic mesoderm cells do not rely on Rac1 and RhoA for movement, which is consistent with their paucity in actin-rich protrusions.

**Figure 6:**
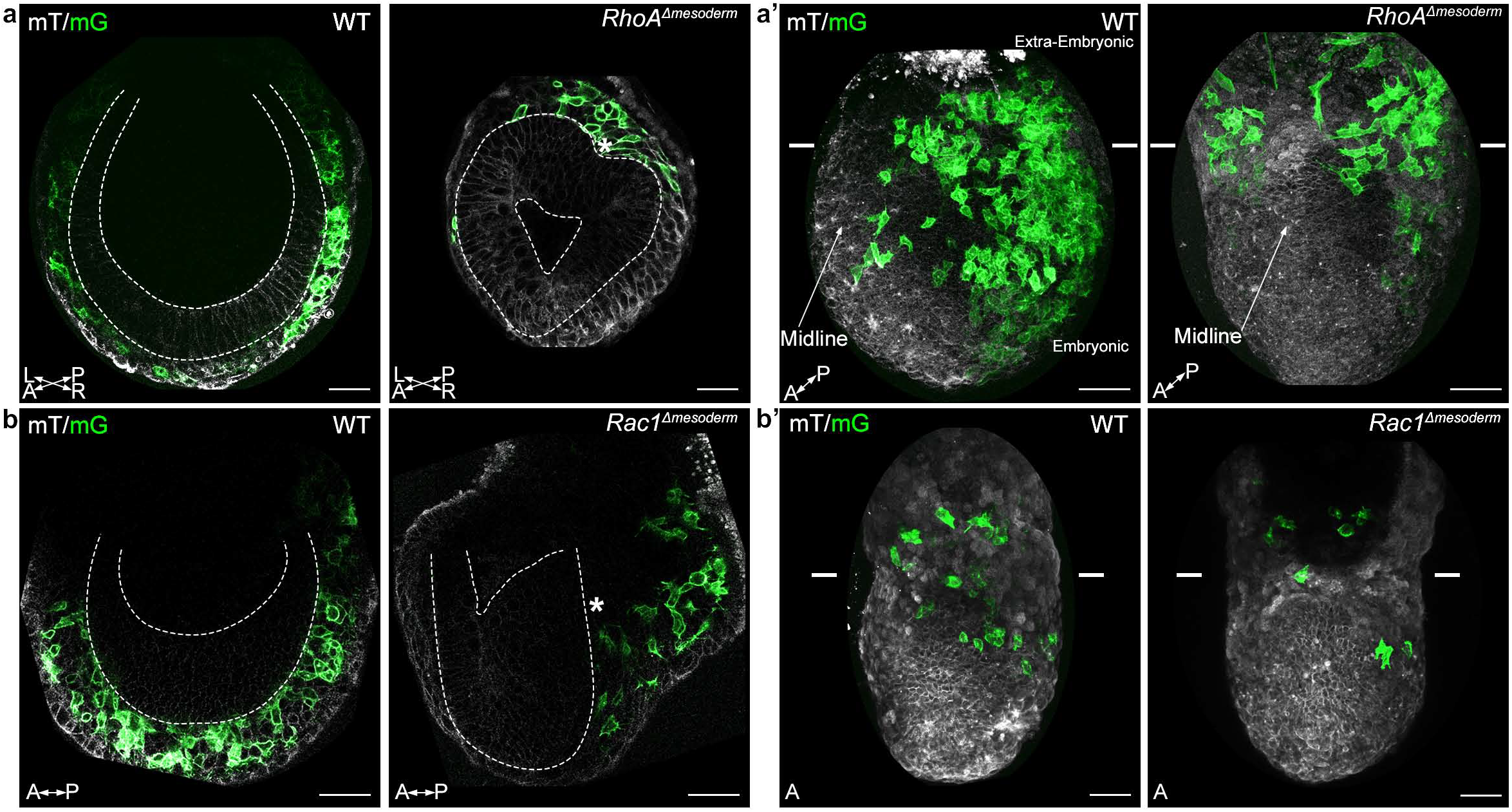
*RhoA and Rac1* mesoderm-specific mutants display impaired migration of embryonic mesoderm. (a) Frontal optical slice of a RhoAL’:.mesoderm embryo and (b) sagittal optical slice of a Rac1L’:.mesoderm embryo highlight accumulation of mesoderm cells next to the primitive streak (*). Dashed lines mark the epiblast. (a’ and b’) Z-projections of stacks from (a’) a RhoAL’:.mesoderm embryo (oblique anterior view, posterior to the right, two-photon) and (b’) a Rac1L’:.mesoderm embryo (anterior view, confocal) show impaired mesoderm migration in embryonic but not extra-embryonic regions. Each mutant is compared to a wild-type littermate in similar orientation. White lines mark the embryonic/extra-embryonic boundary. mG: membrane GFP, in green; mT: membrane dtTomato, in grey. (Scale bar: 50 μm).

## DISCUSSION

Mesoderm cell delamination from the epiblast requires basal membrane disruption, apical constriction, loss of apicobasal polarity, changes in intercellular adhesion, and acquisition of motility (M. Angela Nieto, Huang, Jackson, & Thiery, 2016). The transcriptional network and signaling pathways involved in epithelial-mesenchymal transition are conserved (Ramkumar & Anderson, 2011). However, pre-gastrulation embryo geometry varies widely between species, which has important consequences on interactions between germ layers and mechanical constrains on nascent mesoderm cells (M. L. Williams & Solnica-Krezel, 2017). Live imaging of mouse embryo has allowed recording posterior epiblast rearrangements, as well as cell passage through the primitive streak (Ramkumar et al., 2016; M. Williams et al., 2012). Contrary to the chick embryo, there is no global epiblast movement towards the primitive streak in the mouse. However, cell shape changes, including apical constriction and basal rounding, are similar. Morphological data on mouse mesoderm cells acquired through scanning electron microscopy of whole mount samples (Migeotte et al., 2011), and transmission electron microscopy of embryo sections (Spiegelman & Bennett, 1974) revealed an array of stellate individual cells linked by filopodia containing a lattice of microfilaments.

We took advantage of mosaic labeling of nascent mesoderm to define the dynamics of cell shape changes associated with mesoderm migration. Cells just outside the streak retracted the long apical protrusion and adopted a round shape with numerous filopodia making contacts with adjacent, but also more distant mesoderm cells. In mesodermal wings, cells close to the epiblast were more loosely apposed, and extended fewer filopodia towards its basal membrane, compared to cells adjacent to the visceral endoderm, which were tightly packed and displayed numerous filopodia pointing to the visceral endoderm basal membrane.

Cells travelling in a posterior to anterior direction towards the midline displayed long protrusions, up to twice the cell body size, which extended, retracted, occasionally bifurcated, several times before the cell body itself initiated movement, suggesting an explorative behavior. Remarkably, extension of long protrusions was not limited to the first row of cells. Migration was irregular in time and space, as cells often stopped and tumbled, and displayed meandrous trajectories. After division, cells remained attached by thin protrusions, and followed parallel paths. Contrary to neural crest cells, mesoderm cells did not show contact inhibition of locomotion.

Extra-embryonic mesoderm first accumulates between extra-embryonic ectoderm and visceral endoderm at the posterior side of the embryo, leading to formation of the amniochorionic fold that bulges into the proamniotic cavity (Pereira et al., 2011). This fold expands, and lateral extensions converge at the midline. Accumulation and coalescence of lacunae between extra-embryonic mesoderm cells of the fold generate a large cavity closed distally by the amnion, and proximally by the chorion. At LS stage, extra-embryonic mesoderm forms the allantoic bud, precursor to the umbilical cord, in continuity with the primitive streak (Inman & Downs, 2007). Extra-embryonic mesoderm cells had striking differences in morphology and migration mode, compared to embryonic mesoderm cells. They were larger and more elongated, displayed fewer filopodia, and almost no large protrusions. They migrated at a similar speed, but in a much more tortuous fashion, resulting in little net displacement.

Direction cues could come from cell-matrix contact, homotypic or heterotypic (with epiblast or visceral endoderm) cell-cell interaction, diffuse gradients of morphogens, and/or mechanical constraints. Transcriptome data were compatible with roles for guidance molecules such as Netrin1 and Eph receptors in directing mesoderm migration. *EphA4* was strongly expressed in the PS and mesoderm, particularly in the embryonic region. In *Xenopus*, interaction of EphA4 in mesoderm and Efnb3 in ectoderm allows separating germ layers during gastrulation (Rohani, Parmeggiani, Winklbauer, & Fagotto, 2014). *Epha1* and its ligands *Efna1* and *3* had partially overlapping, but essentially reciprocal compartmentalized expression patterns during gastrulation (Duffy et al., 2006). In addition, we found abundant *Epha1* expression in somites and presomitic mesoderm at E8.5 (not shown). Interestingly, *Epha1* KO mice present a kinked tail (Duffy et al., 2008). The specific and dynamic expression patterns of Epha4, Epha1, and their respective ligands during gastrulation are compatible with roles in germ layers separation, including nascent mesoderm specification and migration. Identification of those potential guidance cues will help design strategies to better understand how mesoderm subpopulations reach their respective destinations.

Visualization and modification of Rho GTPases activity through FRET sensors and photoactivable variants has shed light on their fundamental role during cell migration in *in vivo* contexts, such as migration of fish primordial germ cells (Kardash et al., 2010) or *Drosophila* border cells (Wang, He, Wu, Hahn, & Montell, 2010). Study of a epiblast-specific mutant showed that Rac1 acts upstream of the WAVE complex to promote branching of actin filaments, lamellipodia formation, and migration of nascent mesoderm (Migeotte et al., 2011). Remarkably, extra-embryonic mesoderm cells did not display leading edge protrusions, and *Rac1* and *RhoA* mesoderm-specific mutants were deficient for embryonic, but not extra-embryonic mesoderm migration. In addition, F-actin filaments were more abundant in embryonic mesoderm, reinforcing the hypothesis that they might rely on distinct cytoskeletal rearrangements.

Intermediate filaments are major effectors of cell stiffness, cell-matrix and cell-cell adhesion, as well as individual and collective migration (Pan, Hobbs, & Coulombe, 2013). Members of type I and II keratin families form obligate heterodimers, which assemble into filaments (Loschke, Seltmann, Bouameur, & Magin, 2015). Type II Keratins 7 and 8, and type I Keratins 18 and 19 are the first to be expressed during embryogenesis. Combined Keratins 8/19 and 18/19 deletions cause lethality at E10 attributed to fragility of giant trophoblast cells (Hesse, Franz, Tamai, Taketo, & Magin, 2000). Deletion of the entire type II Keratins cluster results in growth retardation starting at E8.5 (Vijayaraj et al., 2009). Recently, knockdown of Keratin 8 in frog mesendoderm highlighted a role for intermediate filaments in coordinating collectively migrating cells. Keratin-depleted cells were more contractile, displayed misdirected protrusions and large focal adhesions, and exerted higher traction stress (Sonavane et al., 2017). Transcripts for *Keratins 8* and *18*, as well as *Vimentin,* were enriched in extra-embryonic, compared to embryonic mesoderm. While Vimentin was present in all mesoderm, Keratin 8 was only detectable in extra-embryonic mesoderm. An antagonistic relationship between Vimentin intermediate filaments and Rac1-mediated lamellipodia formation has been described (Helfand et al., 2011), and a similar opposition may exist between Rac1 activity and keratin intermediate filaments (Weber, Bjerke, & DeSimone, 2012). Extra-embryonic mesoderm cells’ elongated morphology, paucity in lamellipodia, and lack of directional migration may thus result from their high content in intermediate filaments, and low Rho GTPase activity (Fig. 7). The recent development of a K8-YFP reporter mouse strain for intermediate filaments (Schwarz, Windoffer, Magin, & Leube, 2015), and the availability of reliable Rho GTPases FRET sensors (Spiering & Hodgson, 2011), will be instrumental in dissecting their relationship in mesoderm.

**Figure 7:**
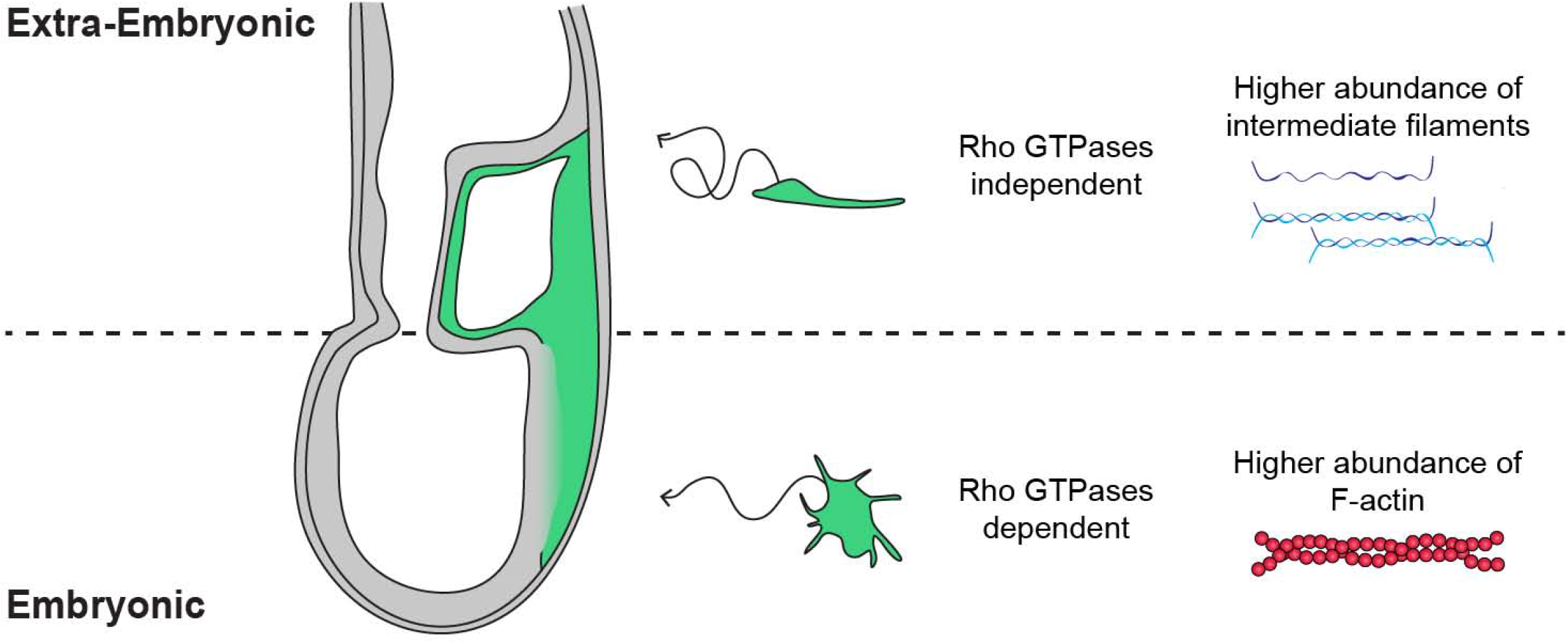
Embryonic and extra-embryonic mesoderm cells show distinct forms, trajectories, Rho GTPases dependency, and cytoskeleton composition during gastrulation in the mouse embryo. In the extra-embryonic region (top), mesoderm cells are stretched, with higher Keratin 8 and Vimentin abundance. Their displacement is convoluted and not dependent on Rho GTPases. In the embryonic region (bottom), mesoderm cells are compact with numerous filopodia, and have higher F-actin abundance. Cells have straighter trajectories and require Rho GTPases. The embryo scheme represents a sagittal section with anterior to the left. Green and grey label mesoderm and other layers, respectively.

The mesoderm germ layer has the particularity to invade both embryonic and extra-embryonic parts of the conceptus, and its migration is important for both fetal morphogenesis and development of extra-embryonic tissues including the placenta. We found that embryonic and extra-embryonic mesoderm populations, both arising by epithelial-mesenchymal transition at the primitive streak, display distinct shape dynamics, migration modes, Rho GTPase dependency, cytoskeletal composition, as well as expression of different sets of guidance, adhesion, and matrix molecules. Landmark experiments in the 1990s showed that the fate of mesoderm cells depends on the time and place at which it emerges from the primitive streak. We have unveiled morphological and behavioral specificities of mesoderm populations through whole embryo live imaging, and provided a molecular framework to understand how cells with distinct fates adapt to, and probably modify, their tridimensional environment.

## ACKNOWLEDGMENTS

We thank the animal house, FACS, and light microscopy (LiMiF) core facilities at the ULB (Erasme Campus), and the Brussels Interuniversity Genomics High Throughput core (www.brightcore.be). We thank M. Martens and J-M. Vanderwinden for confocal imaging help, A. Lefort and F. Libert for RNA sequencing help, C. Brakebusch, A. Gossler, V. Tybulewicz, and L. Machesky for kindly sharing mouse lines. B.S. has been sequentially supported by a fellowship from Erasmus Mundus Phoenix and a fellowship of the FRS/FRIA. N.M. received a fellowship of the FRS/FRIA, as well as support from the "Fonds David et Alice van Buuren" and the "Fondation Jaumotte-Demoulin". W. N. is supported by WELBIO. I.M. is a FNRS research associate and an investigator of WELBIO. WELBIO, the FNRS, and the Fondation Erasme supported this work. The authors declare no financial or non-financial competing interests.

## AUTHOR CONTRIBUTIONS

B.S., N.M., W.N. and I.M. designed the experiments and performed data analysis. B.S., N.M., W.N. and M-L. R. performed experiments. B.S. and W.N. quantified live imaging data, and W.N. wrote the python scripts providing statistical analysis. W.N. drew the schemes. M.D. performed statistical analysis of RNASeq data. I.M. wrote the manuscript with help from B.S. and W.N.

## METHODS

### Mouse strains and genotyping

The *T*-Cre line was obtained from Achim Gossler (Feller et al., 2008), the *Rac1* line from Victor Tybulewicz (Walmsley, 2003), the *RhoA* line from Cord Brakebusch (Jackson et al., 2011), the mTmG (Muzumdar et al., 2007) line from Jax Laboratory, and the LifeAct-GFP line from Laura Machesky (Schachtner et al., 2012). Mice were kept on a CD1 background. Mice colonies were maintained in a certified animal facility in accordance with European guidelines. Experiments were approved by the local ethics committee (CEBEA).

Mouse genomic DNA was isolated from ear biopsies following overnight digestion at 55°C with 1.5% Proteinase K (Quiagen) diluted in Lysis reagent (DirectPCR, Viagen), followed by heat inactivation.

### Embryo culture and live imaging

Embryos were dissected in Dulbecco’s modified Eagles medium (DMEM) F-12 (Gibco) supplemented with 10% FBS and 1% Penicillin-Streptomycin and L-glutamine and 15 mM HEPES. They were then cultured in 50% DMEM-F12 with L-glutamine without phenol red, 50% rat serum (Janvier), at 37°C and 5% CO_2_. Embryos were observed in suspension in individual conical wells (Ibidi) to limit drift, under a Zeiss LSM 780 microscope equipped with C Achroplan 32x/0.85 and LD C Apochromat 40x/1.1 objectives. Stacks were acquired every 20 minutes with 3 μM Z intervals for up to 10 hours. Embryos were cultured for an additional 6 to 12 hours after imaging to check for fitness.

### Antibodies

Antibodies were goat anti-Pecam-1 1:500 (R&D systems); rabbit anti-Podocalyxin 1:200 (EMD Millipore); rat anti-Keratin 8 1:100 (Developmental Studies Hybridoma Bank); rabbit anti-Vimentin 1:200 (abcam). F-actin was visualized using 1.5 U/ml TRITC-Phalloidin (Invitrogen), and nuclei using DAPI (Sigma). Secondary antibodies were anti rabbit Alexa Fluor 488 and 647, anti rat Alexa Fluor 647 (all from Life technologies), and anti goat Alexa Fluor 647 (Jackson).

### Embryo Analysis

Whole-mount *in situ* hybridization was carried out as described in (Eggenschwiler & Anderson, 2000). For *in situ* hybridization on sections, embryos were dissected in PBS and fixed for 30 minutes at 4°C in 4% PFA. They were washed in PBS, embedded directly in OCT (Tissue-Tek), and cryosectioned at 7-10 microns. Slides were re-fixed for 15 minutes on ice in 4% PFA. RNA probes were obtained from ACDBio, and hybridization was performed using the RNAscope 2.5 HD Reagent Kit-RED (ACDBio) according to manufacturer’s instructions. Slides were counterstained with 50% Gill’s Hematoxylin.

For immunofluorescence, embryos were fixed in PBS containing 4% paraformaldehyde (PFA) for 2 hours at 4°C, cryopreserved in 30% sucrose, embedded in OCT and cryosectioned at 7-10 microns. Staining was performed in PBS containing 0.5% Triton X-100 and 1% heat-inactivated horse serum. Sections and whole-mount embryos were imaged on a Zeiss LSM 780 microscope.

### Explant culture and analysis

Primary explant cultures of nascent mesoderm were generated as described in (Burdsal, Damsky, & Pedersen, 1993). Explants were cultured for 24-48 hours in DMEM F-12 supplemented with 10% FBS and 1% Penicillin-Streptomycin and L-glutamine on fibronectin (Sigma) coated glass bottom microwell 35mm dishes with 1.5 cover glass (MatTek). They were fixed for 30 minutes in PBS containing 4% PFA prior to staining. For live imaging, explants were let to adhere for 4-6 hours, and then imaged every 15 minutes for up to 12 hours.

### Image analysis

Images were processed using Arivis Vision4D v2.12.3 (Arivis, Germany). Embryo contours were segmented manually on each Z-slice and time point, and then registered using the drift correction tool of Arivis Vision4D. Embryo rotation was adjusted manually if necessary. We chose embryos where successful registration could be achieved, so that the embryo’s residual slight movements were much smaller than cell displacement. Similarly, we found embryo growth to be negligible compared to cell displacement (data not shown). Cells were then manually segmented on each Z-slice and time point by highlighting cellular membranes using Wacom’s Cintiq 13HD.

Net displacement, path length, speed and angle between two cells were based on the centroid coordinates of segmented cells from Arivis, and calculated by a homemade Python script (Python Software Foundation, https://www.python.org). To extract speed behavior, we interpolated the path length curve and derivated it. The path length over time was closely linear, so we extracted the mean of the speed values. Surface, volume, long/short axis ratio of 2D inner ellipse, and straightness were calculated by Arivis. 2D Z projections of late embryos were used to quantify the filopodia length and density. Filopodia size and density were measured on Icy (de Chaumont et al., 2012) and analyzed using a homemade Python script.

Videos were generated using the StackReg ImageJ plugin (Thevenaz, Ruttimann, & Unser, 1998).

All data are presented as Mean ± SEM. Depending on whether data had a Gaussian distribution or not, we used either the Mann-Whitney-Wilcoxon or the *t*-test. A *p* value <0.05 was considered statistically significant.

### Transcriptome analysis

*T*-Cre; mTmG embryos were collected from different mice, and those at the appropriate stage were pooled. Embryonic and extra-embryonic portions were separated by manually cutting the embryo with finely sharpened forceps. The embryos were digested using 2X Trypsin, and pure GFP+ populations were sorted through flow cytometry (FACSARIA III, BD), directly in extraction buffer. RNA was extracted using the PicoPure kit (ThermoFisher Scientific). RNA quality was checked using a Bioanalyzer 2100 (Agilent technologies). Indexed cDNA libraries were obtained using the Ovation Solo RNA-Seq System (NuGen) following manufacturer recommendation. The multiplexed libraries (18 pM) were loaded on flow cells and sequences were produced using a HiSeq PE Cluster Kit v4 and TruSeq SBS Kit v3-HS from a Hiseq 1500 (Illumina). Paired-end reads were mapped against the mouse reference genome (GRCm38.p4/mm10) using STAR software to generate read alignments for each sample. Annotations Mus_musculus.GRCm38.87.gtf were obtained from ftp.Ensembl.org.

For transcript quantification, all the Reference Sequence (RefSeq) transcript annotations were retrieved from the UCSC genome browser database (mm10). Transcripts were quantified using the featureCounts (Liao, Smyth, & Shi, 2014) software tool using the UCSC RefSeq gene annotations (exons only, gene as meta features). Normalized expression levels were estimated using the EdgeR rpm function and converted to log2 FPKM (fragments per kilobase of exon per million mapped reads) after resetting low FPKMs to 1 to remove background effect. Differential analysis was performed using the edgeR method (quasi-likelihood tests) (McCarthy, Chen, & Smyth, 2012). The edgeR model was constructed using a double pairwise comparison between embryonic mesoderm versus extra-embryonic mesoderm at two different time points (MS and LS). First, the count data were fitted to a quasi-likelihood negative binomial generalized log-linear model using the R glmQLFit method. To identify differentially expressed genes, null hypothesis EM_E7.0 == EEM_E7.0 and EM_E7.25 == EEM_E7.25 were tested using the empirical Bayes quasi-likelihood F-tests (glmQLFTest method) applied to the fitted data. The F-test P-values were then corrected for multi-testing using the Benjamini-Hochberg p-value adjustment method. Transcripts with a greater than background level of expression (mean log2 count per million > 0), an absolute fold change > 2, and a low false discovery rate (FDR < 0.05) were considered as differentially expressed.

The sample visualization map was produced by applying the t-Distributed Stochastic Neighbor Embedding (tSNE) dimensionality reduction method (Van Der Maaten & Hinton, 2008) to log2 FPKM expression levels (all transcripts). The R tSNE method from ’Rtsne’ library was applied without performing the initial PCA reduction and by setting the perplexity parameter to 2. The heatmap was produced using the R heatmap.2 methods using the brewer.pal color palette. GO analysis was performed using the DAVID software (Huang, Sherman, & Lempicki, 2009).

## LEGENDS FOR VIDEOS

**Supplementary Video 1: Mesoderm cells migrating towards extra-embryonic and embryonic regions.** Z-projections of confocal stacks from a *T*-Cre; mTmG embryo dissected at E6.75 (Early Streak stage) and imaged for 320 min. Mesoderm cells express membrane GFP (green); all other cells express membrane dtTomato (red). Anterior oblique orientation with posterior to the right (scale bar: 50 μm).

**Supplementary Video 2: "Trial and error" trajectories.** Z-projections of confocal stacks from a *T*-Cre; mTmG embryo dissected at E6.75 (Mid Streak stage) and imaged for 260 min. Mesoderm cells express membrane GFP (green); all other cells express membrane dtTomato (red). Anterior oblique orientation with posterior to the right (scale bar: 50 μm).

**Supplementary Videos 3 and 4: Tracking mesoderm migration.** 3D snapshots of confocal stacks from *T*-Cre; mTmG embryos dissected at E6.75 (Early Streak stage) and imaged for 180 min and 160 min, with manually highlighted cells tracked throughout the time lapse. Videos show highlighted cells first, then the original images (membrane GFP, in green) in a looping fashion for comparison. All other cells express membrane dtTomato (grey). Lateral orientation with anterior to the left (scale bar: 50 μm).

**Supplementary Video 5: Extra-embryonic mesoderm migration is characterized by low net displacement.** Z-projection of confocal stack from a *T*-Cre; mTmG embryo dissected at E6.75 (Early Streak stage) and imaged for 860 min cropped to show extra-embryonic mesoderm cells. Mesoderm cells express membrane GFP (green) (scale bar: 10 μm).

**Supplementary Video 6: Mesoderm extends filopodia towards epiblast and visceral endoderm.** Two-Photon stack of a *T*-Cre; mTmG embryo at Late Streak stage.

The stack progresses from anterior to posterior. Mesoderm cells express membrane GFP (green); all other cells express membrane dtTomato (grey) (scale bar: 50 μm).

**Supplementary Video 7: Searching behaviour.** Z-projection of two-photon stack from a *T*-Cre; mTmG embryo at Mid Streak stage imaged for 100 min. Arrow points at embryonic mesoderm cell. Mesoderm cells express membrane GFP (green); all other cells express membrane dtTomato (grey) (scale bar: 50 μm).

**Supplementary Video 8: LifeAct-GFP expression is higher in embryonic mesoderm.** 3D snapshots of two-photon stacks from a *T*-Cre; LifeAct-GFP embryo dissected at E7.25 (Late Streak stage) and imaged for 295 min. Images were processed with the ZEN blue denoise function. LifeAct-GFP (in green) highlights F-actin. The bright specks in extra-embryonic on the right side are debris. Lateral orientation with anterior to the left (3D scale bar: 50 μm).

**Supplementary Video 9 and 10: *RhoA*Δ*mesoderm* explants undergo compaction before cell migration compared to wild type.** Z-projection of confocal stack of mesoderm explants from *T*-Cre; mTmG (9) and *T*-Cre; mTmG; *RhoA* fl/- (10) embryos dissected at E7.5 (Late Streak stage). Explants were imaged for 750 min every 15 min.

**Supplementary Video 11: *RhoA*Δ*mesoderm* embryos display an accumulation of mesoderm near the primitive streak.** Two-photon Z stack of a *T*-Cre; mTmG; *RhoA* fl/- embryo at Late Streak stage. Mesoderm cells express membrane GFP (green); all other cells express membrane dtTomato (grey). Anterior oblique orientation with anterior to the left (scale bar: 50 μm).

**Supplementary Video 12: Mesoderm migration tracking in a *RhoAΔmesoderm* embryo.** 3D snapshots of two-photon stacks from a *T*-Cre; mTmG; *RhoA* fl/- embryo dissected at E7.25 (Mid Streak stage) and imaged using 2-Photon confocal microscopy for 120 min showing highlighted cells, which are tracked throughout the time lapse. The video shows the highlighted cells first, then the original images (Membrane GFP, in green) in a looping fashion for comparison. All other cells express membrane dtTomato (grey). Anterior oblique orientation with anterior to the left (scale bar: 50 μm).

**Supplementary Video 13: Mesoderm migration tracking in *Rac1Δmesoderm* embryos.** 3D snapshots of two-photon stacks from a *T*-Cre; mTmG; *Rac1* fl/- embryo dissected at E7.25 (Mid Streak stage) and imaged for 80 min showing highlighted cells, which are tracked throughout the time lapse. The video shows the highlighted cells first, then the original images (Membrane GFP, in green) in a looping fashion for comparison. All other cells express membrane dtTomato (grey). Lateral orientation with anterior to the left (scale bar: 50 μm).

## LEGENDS FOR SOURCE DATA

**Source Data 1. Embryonic and extra-embryonic mesoderm cells tracking:** List detailing individual cells tracking, volume and surface measurement results.

**Source Data 2. Embryonic and extra embryonic mesoderm shape measurements**

**Source Data 3. Embryonic and extra embryonic mesoderm cells filopodia:**
Filopodia number/cell and filopodia length measurements.

**Source Data 4. Mesoderm cells filopodia extended towards Visceral Endoderm and Epiblast.**

**Source Data 5. Quantification of mesoderm cellular division.**

**Source Data 6. Quantification of mesoderm cellular collision.**

**Source Data 7. Description and quality control of samples used for RNA-seq.** EM: Embryonic mesoderm; EEM: Extra-Embryonic mesoderm; EC: Embryonic control; EEC: Extra-Embryonic control.

**Source Data 8: Expression Levels.** Table containing expression levels in log2 FPKM computed using the rpkm edgeR method.

**Source Data 9: Ranked list of differential expression.** Column 1: gene name, Column 2 log2 Fold change between EM_E7.0 and EEM_E7.0, Column 3 log2 Fold change between EM_E7.25 and EEM_E7.25, Column 4 log2 Count Per Million, Column 5 F-test value, Column 6 F-test pvalue, Column 7 F-test FDR (Benjamini-Hochberg).

**Supplementary Figure 1:**
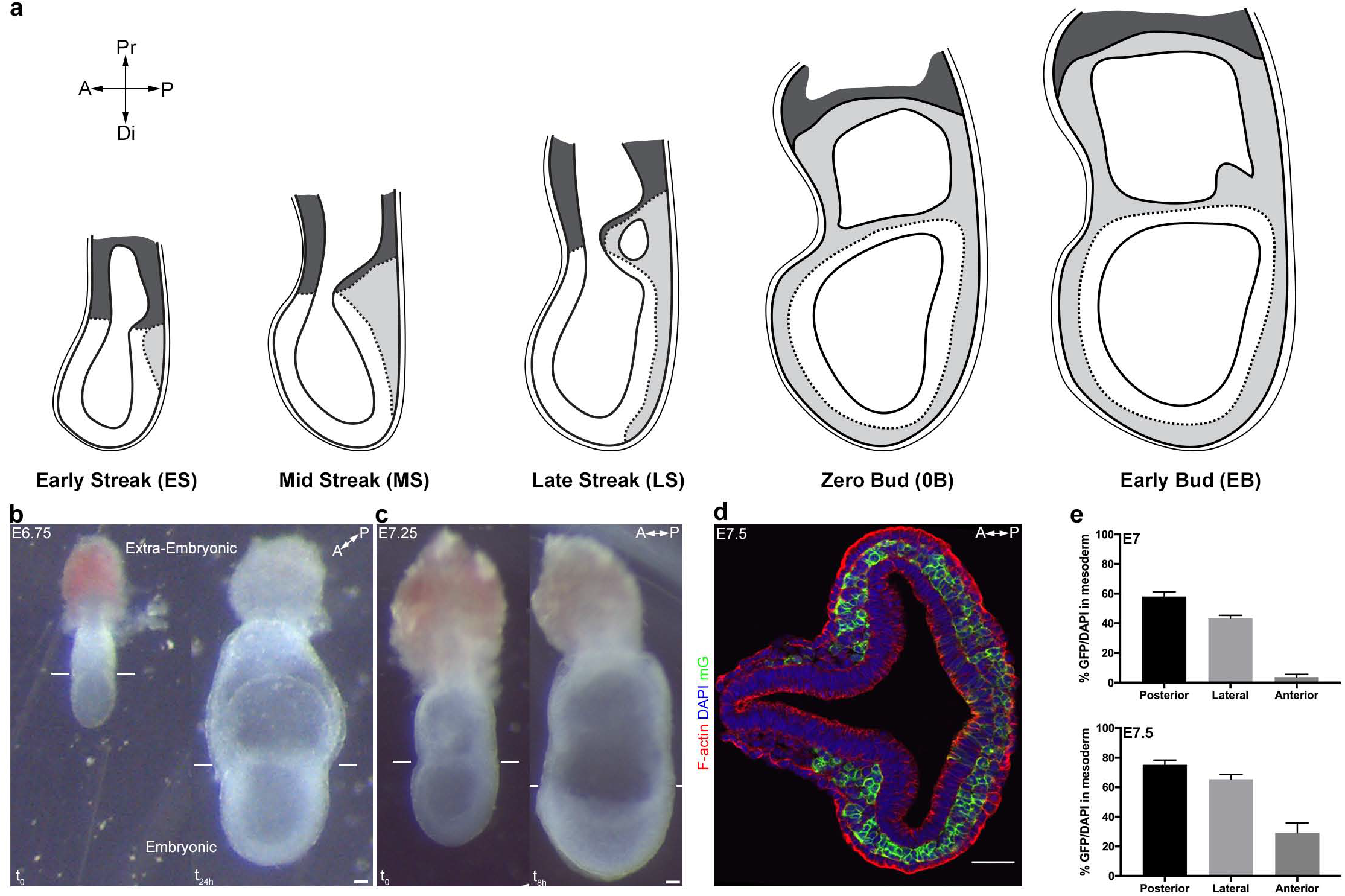
Live imaging of *Brachyury* (*T*)-Cre; mTmG embryos. (a) Sketches showing mouse embryo anatomical landmarks used for staging between Early Streak and Early Bud stages. (b) *Ex vivo* growth of an embryo dissected at E6.75 (Early Streak stage, left panel), after imaging for 12 hours using confocal imaging plus 12 hours of subsequent culture (Zero Bud stage, right panel). (c) *Ex vivo* growth of an embryo dissected at E7.25 (Late Streak stage, left panel), after 8 hours two-photon imaging (Early Bud stage, right panel). (d) Transverse section at primitive streak level of a E7.5 (Late Streak stage) T-Cre; mTmG embryo stained for F-actin (Phalloidin, red) and nuclei (DAPI, blue) (scale bar: 50 μm). (e) Quantification of mosaicism as the ratio of GFP+ cells/all cells in the mesoderm layer (identified anatomically between the epiblast and the visceral endoderm) in anterior, posterior and lateral quadrants at different stages of gastrulation: E7 (Mid Streak stage) and E7.5 (Early Bud stage) (mean ± SEM, n=5 embryos for each stage). A: Anterior, P: Posterior, Pr: Proximal, Di: Distal, mGFP: membrane GFP, in green.

**Supplementary Figure 2:**
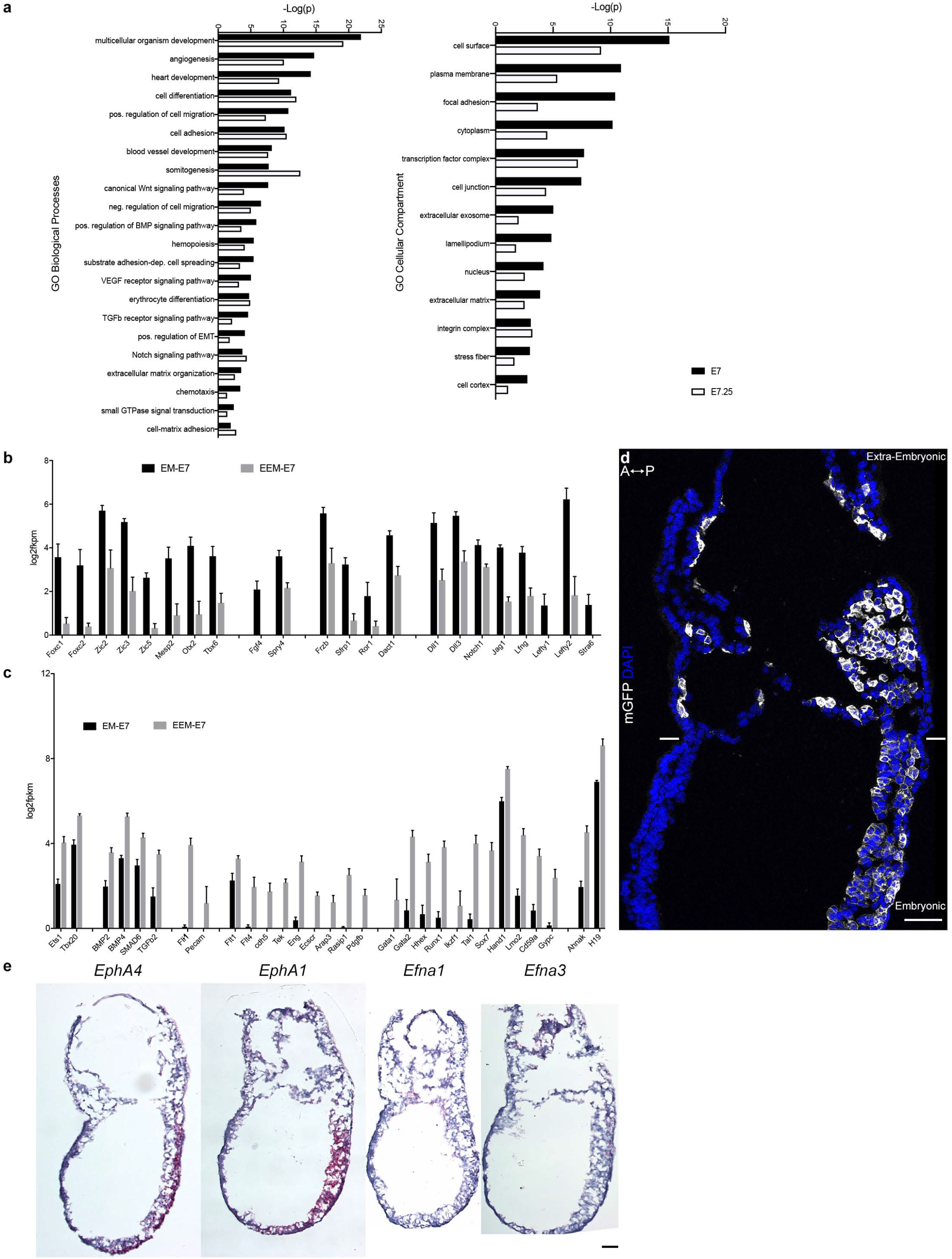
Embryonic and extra-embryonic mesoderm transcriptome in the mouse embryo. (a) Gene ontology enrichment of gene clusters related to biological processes and cellular compartments that are differentially expressed between embryonic and extra-embryonic mesoderm with the higher statistical significance at E? and E?.25. (b,c) Selection of genes known to have higher expression in (b) extra-embryonic or (c) embryonic mesoderm, represented as mean± SEM of log2 fpkm at E? (n=4 biological replicates, p<0.001), with embryonic in black and extra-embryonic in grey. EM: Embryonic mesoderm and EEM: Extra-Embryonic mesoderm. Data can be found in Source Data 8 and 9. (d) Optical slice of sagittal section (anterior to the left) from Early Bud stage embryo with membrane GFP in grey and nuclei in blue (DAPI) (Scale bar: 50 μm) (e) *In situ* hybridization of sagittal sections (anterior to the left) from Zero to Early Bud stages embryo highlighting transcripts from *EphA4, EphA1, Efna1* and *Efna3,* represented by red dots (Scale bar: 100 μm).

**Supplementary Figure 3:**
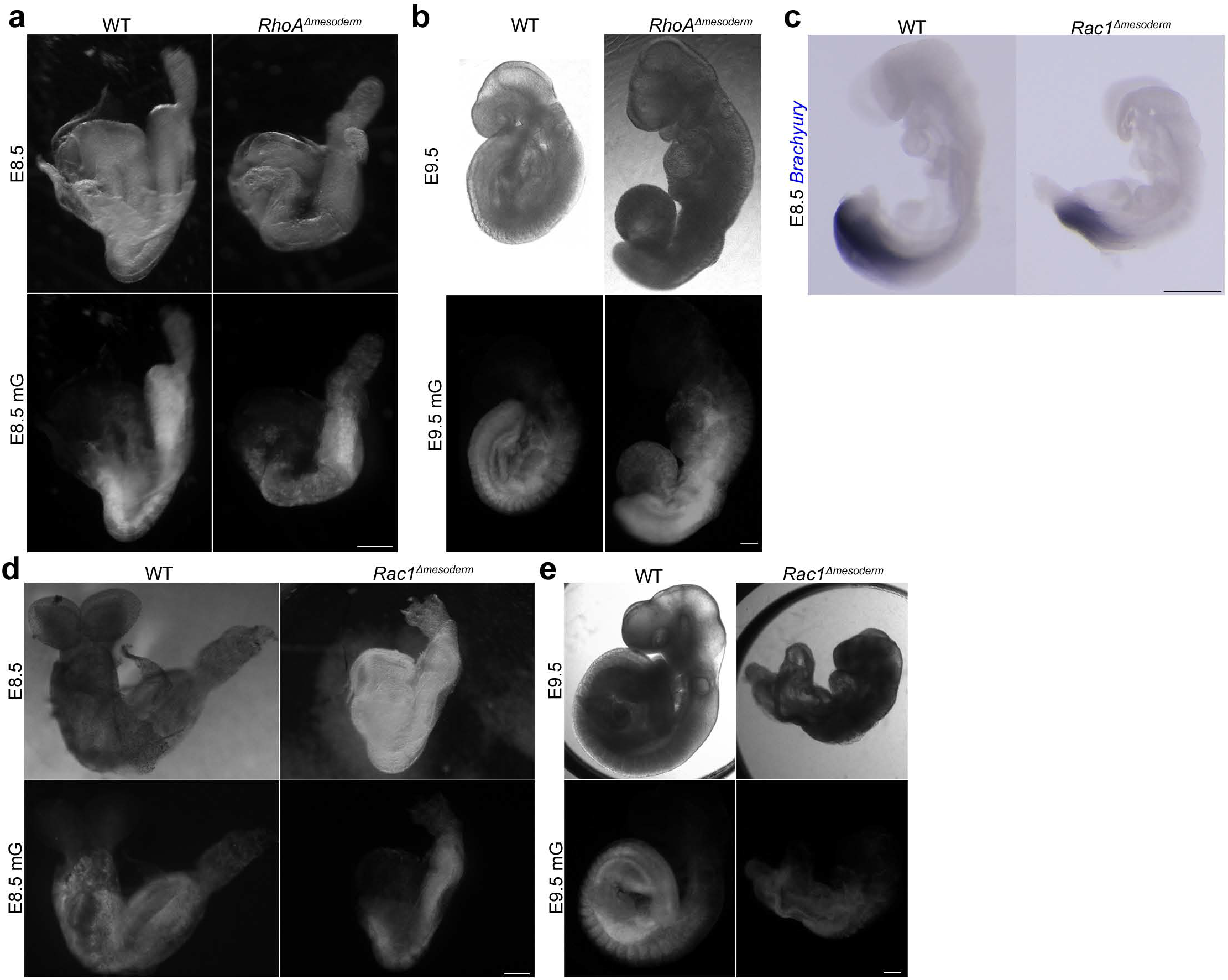
Phenotypes of *Rac1 and RhoA* mesoderm-specific mutants post gastrulation. (a, b) Bright field (top) and mGFP epifluorescence (bottom) images of RhoALlmesoderm embryos at E8.5 and E9.5 (c) *In situ* h ybridization for *Brachyury* at E8.5 of *Rae 1*Llmesoderm embryo showing defect in presomitic mesoderm. (d, e) Bright field (top) and mGFP epifluorescence (bottom) images of Rac1Llmesoderm embryos at E8.5 and E9.5. All mutants are compared to a wild-type littermate. mGFP: membrane GFP, in grey. (Scale bars: 200 μm)

**Supplementary Figure 4:**
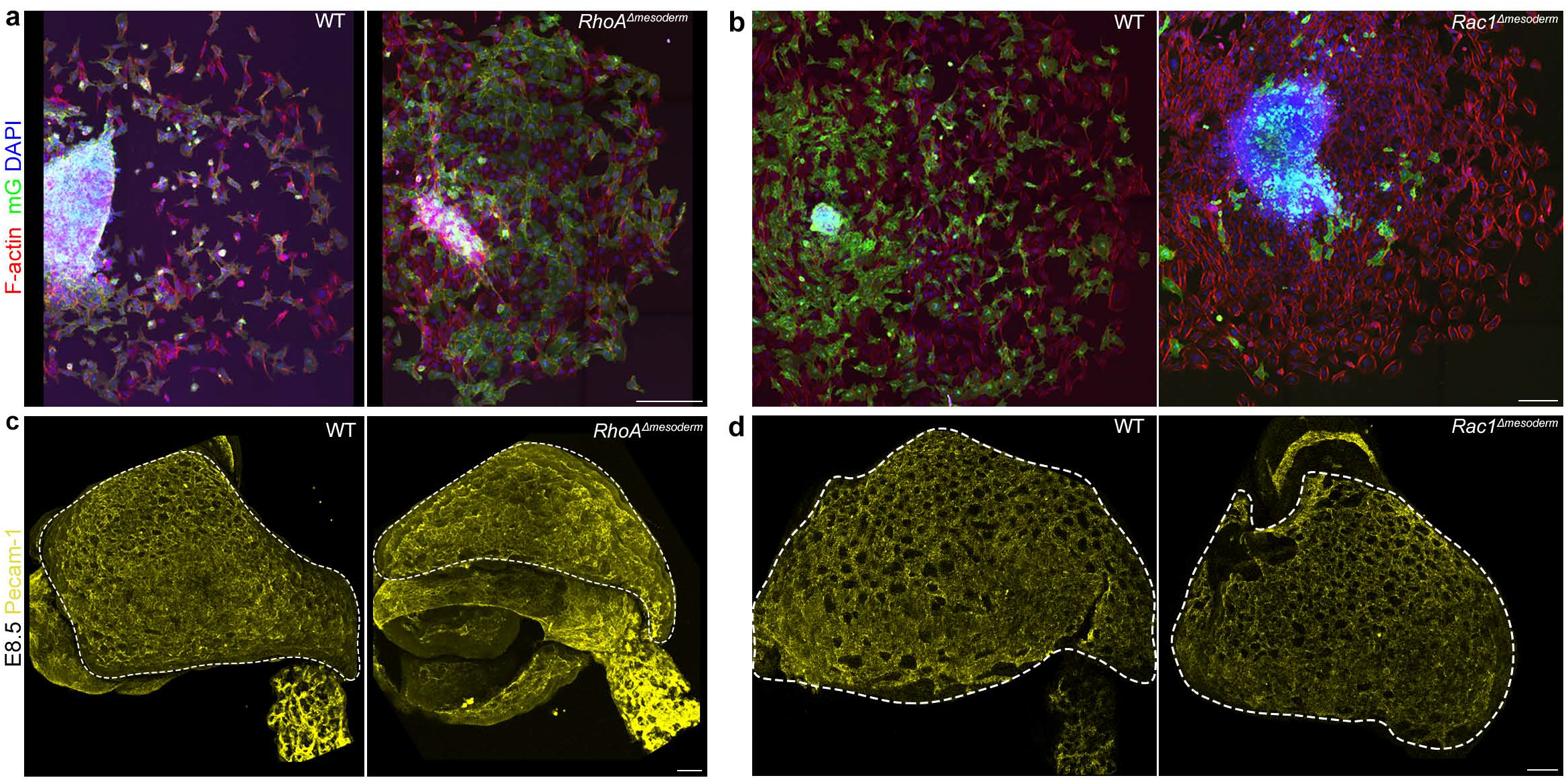
Embryonic and extra-embryonic mesoderm cellular details in *RhoA* and *Rac1* mesoderm-deleted embryos. (a, b) Embryonic mesoderm explants from (a) mTmG; RhoAt:.mesoderm, and (b) mTmG; Raclt:.mesoderm Mid/Late Streak embryos cultured on fibronectin for (a) 48h and (b) 30h, stained for F-actin (Phalloidin, in red) and nuclei (DAPI, in blue). mG: membrane GFP, in green. (N=3 for *Rac1,* 4 for *RhoA;* similar phenotype in 2/3 mutants for *Rac1* and 3/4 mutants for *RhoA).* (Scale bars: 200μm for a, and 50μm for b). (c, d) Whole-mount E8.5 (c) RhoAt:.mesoderm and (d) Raclt:.mesoderm embryos stained for Pecam-1 (in yellow). Dashed lines mark the yolk sack. (N=2 for *RhoA,* 8 for *Rac1).* (Scale bars: 100μm). All mutants are compared to a wild-type littermate.

